# An artifact of recombinatorial cloning challenges established beliefs of plasmid co-transformation, selection, and maintenance

**DOI:** 10.1101/2025.01.14.633004

**Authors:** Courtney L. Geer, J. Michael Charette

**Affiliations:** Department of Chemistry, Brandon University, Brandon, Manitoba, Canada; Children’s Hospital Research Institute of Manitoba, Winnipeg, Manitoba, Canada; Paul Albrechtsen Research Institute, CancerCare Manitoba, Winnipeg, Manitoba, Canada

## Abstract

Gateway cloning is an easy, efficient, accurate, and versatile cloning strategy. During Expression clone validation, we sometimes see an additional band co-migrating with the pDONR backbone. We show that this “mystery” band is not an artifact of aberrant recombination but instead originates from a co-transformation event. We find that the unselected pDONR plasmid is co-transformed into *E. coli* with the desired Expression vector in 9–29% of colonies and is maintained without antibiotic selection, despite plasmid incompatibility. We propose an easy strategy to screen for co-transformants. Our results challenge accepted beliefs of bacterial plasmid transformation, selection, and maintenance.

## Description

### Traditional cloning in comparison to Gateway cloning

The advent of Gateway recombinational cloning in the late 1990’s revolutionised cloning technology due to its greatly improved speed, efficacy, accuracy, and flexibility compared to traditional methods using restriction endonucleases and DNA ligation. Gateway cloning was instrumental to the development of high-throughput systems biology studies requiring the generation of hundreds to thousands of clones. This included the creation of proteome-scale clone libraries such as the yeast Movable Open Reading Frame (MORF) collection (Gelperin et al., 2005), the Human ORFeome (Rual et al., 2004), the *Xenopus* ORFeome (Grant et al., 2015), *Arabidopsis* cDNA libraries (Bürkle et al., 2005), the *Vibrio cholerae* FLEXGene clones (Rolfs et al., 2008), and countless other organismal ORF clone collections. Gateway cloning enables the easy movement of these ORFs into different expression vectors for their functional characterization. The scalability of Gateway cloning facilitates this at the low (Charette & Baserga, 2010), medium (McCann et al., 2015; Vincent et al., 2018), and high-throughput (Rolland et al., 2014; Yu et al., 2008) levels. While expensive relative to traditional restriction enzyme and ligation-based cloning, Gateway cloning’s high success rate makes it a cloning method of choice for primarily undergraduate research labs and other labs that may lack in-house cloning expertise.

### Fundamentals of Gateway recombinational cloning

Based on the site-specific recombination reactions of bacteriophage lambda (λ) with the *E. coli* genome, Gateway cloning facilitates the movement of DNA inserts between cloning vectors while maintaining reading frame and orientation (Hartley et al., 2000; Landy, 1989). Specifically, attachment (*att*) sequences borrowed from the *E. coli* bacterial chromosome (*att*B) and the lambda phage genome (*att*P) allow for the integration and excision of DNA inserts into and out of Gateway vectors – a process requiring only two different enzyme mixes (Hartley et al., 2000; Katzen, 2007). First, the Gateway BP clonase enzyme mix, consisting of Integrase (Int) and Integration Host Factor (IFH), recombines *att*B and *att*P sites to produce *att*L (left) sites (Suppl. Fig. 1a). Typically, DNA fragments or amplified PCR products flanked on either side by *att*B sites are recombined with a Donor vector containing *att*P sites – which flank both sides of a Gateway cassette (described below). This generates an Entry clone containing the PCR product, now flanked by *att*L sites, and a discarded fragment consisting of the excised Gateway cassette. Entry clones are non-expressing in that they serve only to house the insert for later recombination reactions. The second reaction involves recombination of the *att*L sites in the Entry clone with the *att*R (right) sites of a Destination vector also containing a Gateway cassette. This recombination is catalysed by the LR clonase enzyme mix, consisting of Excisionase (Xis), which excises the DNA insert from the Entry clone, and Int and IHF, which recombine it with the Destination vector, resulting in an Expression vector and a second discarded Gateway cassette (Suppl. Fig. 1b) (Hartley et al., 2000; Katzen, 2007). Because the BP and LR enzymes recognize only their corresponding sequence sites, each recombination reaction is incredibly accurate. As well, the orientation of the DNA insert is maintained using a set of *att* site subtypes (*e*.*g*., *att*B1 and *att*P1) that exclusively recombine with others of the same subtype (Reece-Hoyes & Walhout, 2018).

**Figure 1.**
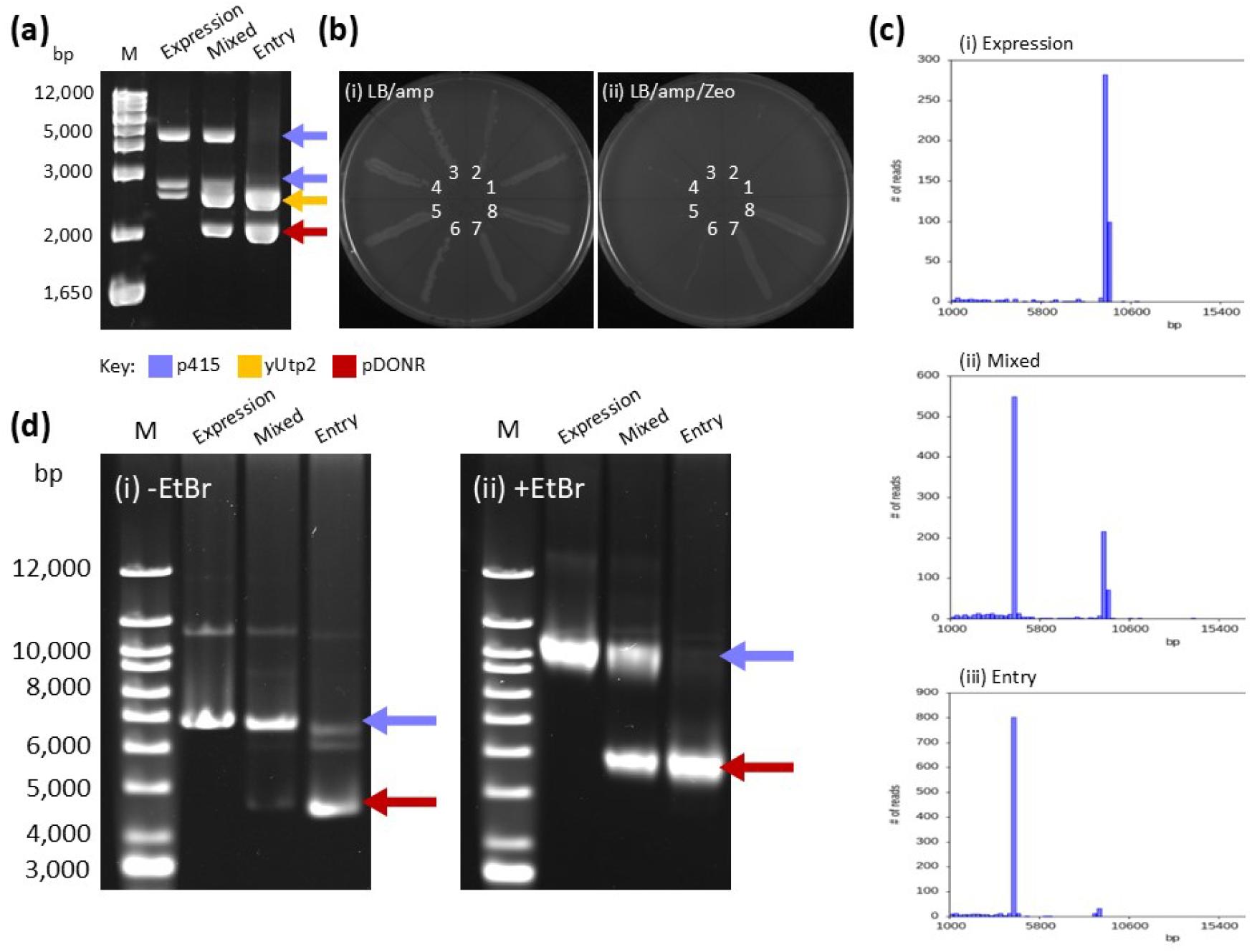
Gateway Entry and Expression vectors are sometimes co-transformed into *E. coli*. **(a)** Gel electrophoresis of *Bsr*GI digests of p415GPD-3xHA-GW-yUtp2 Expression vector clones 1 (Expression Lane; showing ampicillin (amp) resistance) and 7 (Mixed Lane; showing Zeocin (Zeo) and amp resistance) from (1b) and the pDONR-yUtp2 Entry vector (Entry Lane). Sizes of the 1 Kb+ molecular weight ladder (Lane M) are shown to the left. Bands in the Expression Lane at ∼4500 and ∼2700 base pairs (bp) correspond to the p415GPD-3xHA-GW backbone (blue arrows), which produces bands of 4199 and 2570 bps along with a band at 180 bp (not seen). The band at ∼ 2500 bp corresponds to the 2433 bp yUtp2 gene sequence plus the flanking Gateway *att*R sequences (yellow arrow) which add up to 2466 bp. The 4199 and 2570 bp p415GPD-3xHA-GW backbone bands, along with the 2466 bp yUtp2 band (with *att*R sequences), are also present in the Mixed Lane, as is an additional band at ∼2000 bp co-migrating with the 2043 bp pDONR backbone (red arrow). A band of the same size (2043 bp; pDONR backbone) is also seen in the Entry Lane, as well as a second band at ∼2500 bp, corresponding to the 2466 bp yUtp2 sequence plus *att*R sequences. **(b)** Transformants 1-8 of the LR cloning reaction that generated the p415GPD-3xHA-GW-yUtp2 Expression vector were dashed on LB/amp (i) and LB/amp/Zeo (ii) plates. Clones 1-6 only show growth on the LB/amp (i) plate, while clones 7 and 8 demonstrate both amp and Zeo resistance. Plasmid DNA from clones 1 and 7 was analyzed by *Bsr*GI digestion shown in (1a). **(c)** Plasmid population histograms showing (i) a single peak at ∼ 9500 bp corresponding to the p415GPD-3xHA-GW-yUtp2 Expression vector, (ii) two distinct plasmid populations consistent with the sizes of the Expression (∼9500 bp) and Entry vectors (∼4500 bp), and (iii) a distinct peak at ∼ 4500 bp corresponding to the pDONR-yUtp2 Entry vector (with a small peak at ∼ 9000 bp consistent with a plasmid dimer of the same vector). **(d)** Gel electrophoresis of the undigested plasmid DNAs used in 1a and 1c resolved in the absence (i) and presence (ii) of ethidium bromide (EtBr). Sizes of the 1 Kb+ molecular weight ladder (Lane M) are shown to the left on both gels. Bands at ∼ 7000 bp in the Expression and Mixed lanes in the gel without EtBr (i) correspond to the 9415 bp p415GPD-3xHA-GW- yUtp2 plasmid (blue arrow), with the bands at ∼ 4500 bp in the Mixed and Entry lanes matching the size of pDONR-yUtp2 (red arrow) at 4509 bp. The gel containing samples resolved in the presence of EtBr (ii) exhibits bands at ∼ 10 Kbp in the Expression and Mixed lanes, indicating the presence of p415GPD-3xHA-GW-yUtp2 (blue arrow), as well as bands at ∼5500 bp in the Mixed and Entry lanes corresponding to pDONR-yUtp2 (red arrow). Other faint bands may correspond to plasmid topoisomers.

Gateway cloning uses both positive and negative selection. The selection of transformed *E. coli* cells is done as per conventional cloning. The elimination of unrecombined plasmids is made possible through the retention of a Gateway cassette in both Donor and Destination vectors which contains both the toxic *ccdB* gene and a chloramphenicol (Cm^R^) resistance gene (Suppl. Fig. 1). The *ccdB* gene enables negative selection of unrecombined plasmids through the production of a toxic CcdB protein that kills most cloning *E. coli* strains (including DH5α) (Reece-Hoyes & Walhout, 2018). Chloramphenicol resistance is used to propagate and maintain the Gateway cassette within Donor and Destination vectors in cells tolerant to CcdB protein toxicity, such as the *E. coli* DB3.1 strain (Villefranc et al., 2007).

The validation of Gateway clones takes advantage of *Bsr*GI restriction sites in the *att* sequences. Following the isolation of Entry clone plasmid DNA, *Bsr*GI digestion liberates the insert from the vector backbone. Electrophoresis should reveal an insert of the expected size (though possibly containing internal *Bsr*GI sites) along with the vector backbone (both pDONR-Zeocin (Zeo) and pDONR221 Entry vectors contain no *Bsr*GI sites in their backbone). Such putative Entry clones are further validated by DNA sequencing. Expression clones are similarly validated by *Bsr*GI digestion. However, upon identification of an insert of the expected size, the plasmid clone is typically not re-sequenced due to the accuracy of the Gateway recombination reaction (though it is often validated at the protein expression level).

### Bacterial co-transformation

Most researchers accept the idea that transformation involves the movement of a single plasmid molecule into a single bacterial cell (excluding subsequent transformations). This (*i*.*e*., one plasmid into one cell) is thought to be ensured by plasmid incompatibility and antibiotic selection (Sambrook & Russell, 2001; Velappan et al., 2007). Incompatible plasmids have the same origin of replication and are therefore assumed to compete for shared replication factors, leading to the dominance of the plasmid possessing growth advantages and the eventual loss of the second plasmid (Sambrook & Russell, 2001; Velappan et al., 2007). Similarly, antibiotic resistance markers have been used as a means of selecting for and identifying transformed plasmids. For example, it has long been thought that different plasmids encoding the same antibiotic resistance gene are rarely simultaneously transformed (Goldsmith et al., 2007). Following transformation, antibiotic markers are also used as a means of conferring selective advantage to a plasmid of interest, allowing it to be maintained within the cell in the presence of the corresponding antibiotic (Sambrook & Russell, 2001). However, in the absence of the antibiotic, plasmids carrying the unselected antibiotic resistance gene no longer provide a selective advantage and are thus assumed to be lost over time. This is often attributed to the metabolic cost of replicating plasmid DNA that confers no selective advantage (Carroll & Wong, 2018; Vogwill & MacLean, 2015).

The process of co-transformation is the simultaneous transformation of a single bacterial cell by two or more different plasmids. Its assumed rarity has been passed down through laboratory lore as more myth than fact, as it has been thought to be prevented by plasmid incompatibility and antibiotic selection. Unexpectedly, studies by Goldsmith et al. (2007), Velappan et al. (2007), and more recently by Tomoiaga et al. (2022) have demonstrated that the phenomenon may be more common than originally believed. Despite accepted beliefs about plasmid incompatibility, plasmid selection and maintenance as it is facilitated by antibiotic resistance, and the rarity of simultaneous co-transformation of multiple different plasmids, studies using plasmids with identical origins of replication have observed the co-transformation of up to eleven different plasmids into the same *E. coli* cell (Goldsmith et al., 2007; Tomoiaga et al., 2022; Velappan et al., 2007). This includes the co-transformation of two plasmids with the same origin of replication but different antibiotic resistance genes (while selecting for each antibiotic separately and simultaneously for both antibiotics (Goldsmith et al., 2007; Velappan et al., 2007)) and that of nearly identical plasmids each expressing one of several different fluorescent proteins (Tomoiaga et al., 2022).

In our routine cloning, we observe that an additional band is sometimes seen in Gateway cloning Expression vectors during *Bsr*GI restriction digestion validation. This “mystery” band co-migrates with the pDONR Entry backbone, which initially led us to suspect that an aberrant recombination event had occurred during the LR reaction. However, Oxford Nanopore sequencing surprisingly shows that unrecombined pDONR Entry plasmid is instead co-transformed and maintained in the absence of selection alongside the desired Expression vector. To our knowledge, this is the first recorded instance of co-transformation occurring with Gateway cloning. Taken together, our results challenge long-held beliefs of single-plasmid transformation, clonality, antibiotic selection, and maintenance of laboratory plasmids in *E. coli*.

### Plasmid co-transformation occurs during routine Gateway cloning

Following the generation of all new Gateway Expression vectors, our laboratory performs diagnostic *Bsr*GI restriction digestion validation of candidate clones. During this standard procedure, we observe an unexpected ∼2 Kbp band in the digests of some p415GPD-3xHA-GW-yUtp2 Expression vector transformants (a yeast expression plasmid). This additional band co-migrates with the pDONR Entry plasmid backbone (Fig. 1a). Based on the size of this band – corresponding to that of the Entry vector’s backbone – we initially suspected that an aberrant LR recombination event had occurred, resulting in a failure to recombine the Destination vector’s discarded Gateway cassette with the Entry vector backbone. If so, this might result in the Entry pDONR backbone (here, Zeocin) being integrated into the Expression vector (here, ampicillin, or amp), yielding a concatenated plasmid. To test this theory, we screened several *E. coli* transformants (transformed with the products of the LR reaction) from the original LB/amp transformation plate for growth on LB/amp and on LB/amp/Zeo plates (Fig. 1b). Cells possessing the suspected concatenated plasmid were expected to carry the genes for both Zeocin and ampicillin resistance, which would allow for their growth on both antibiotics. We found that two of the eight clones (Fig. 1b (ii), clones 7 and 8) screened displayed both Zeocin and ampicillin resistance. The remaining six clones only grew on ampicillin, as expected.

To further investigate the clones growing on both Zeocin and ampicillin, we took advantage of the recently developed Oxford Nanopore whole plasmid sequencing strategy. We also sequenced plasmid DNA from colonies only exhibiting ampicillin resistance (likely the p415GPD-3xHA-GW-yUtp2 Expression vector) along with the pDONR-yUtp2 Entry vector. To our surprise, Oxford Nanopore sequencing of plasmid DNA extracted from a single bacterial colony resistant to both Zeocin and ampicillin (Fig. 1b, clone 7, cultured overnight in liquid LB/amp (without Zeo) for the plasmid miniprep and Fig. 1a, Mixed Lane) revealed the presence of two plasmid populations, rather than a single large concatenated plasmid (Fig. 1c (ii) and Suppl. Fig. 2a and b). These plasmids of ∼4.5 and ∼9.5 Kbp in size correspond to the anticipated sizes of the Entry plasmid pDONR-yUtp2 and of the Expression vector p415GPD-3xHA-GW-yUtp2. Sequence analysis further confirms this observation, as the ∼4.5 Kbp species corresponds to the 4509 bp pDONR-yUtp2 Entry vector (pDONR-Zeo encoding the *S. cerevisiae* YPH499 NOP14/UTP2 gene). Similarly, the ∼9.5 Kbp species corresponds to the 9415 bp p415GPD-3xHA-GW-yUtp2 Expression vector (p415GPD-3xHA-GW-*att*R (McCann et al., 2015) with the yeast YPH499 NOP14/UTP2). Oxford Nanopore sequencing of plasmid DNA extracted from a bacterial colony that is only ampicillin resistant (Zeo sensitive; Fig. 1b, clone 1 and Fig. 1a, Expression Lane) only contained one plasmid species (Fig. 1c (i) and Suppl. Fig. 2b) with a plasmid size of 9415 bp and a sequence that corresponds to that of p415GPD-3xHA-GW-yUtp2. Oxford Nanopore sequencing of pDONR-yUtp2 yielded a predominant plasmid population of 4509 bp (Fig. 1a, Entry Lane, Fig. 1c (iii), and Suppl. Fig. 2a) with a sequence that corresponds to that of pDONR-yUtp2, as well as a small population of a pDONR-yUtp2 plasmid dimer with a size of ∼ 9000 bp. (Because this small population is twice the size of the pDONR-yUtp2 vector, and because it does not match the size of the p415GPD-3xHA-GW-yUtp2 plasmid (9415 bp), we are confident that it is dimerized pDONR-yUtp2 Entry plasmid.)

In addition, no concatenated plasmid sequences (resulting from an aberrant recombination and fusion between pDONR-yUtp2 and p415GPD-3xHA-GW*-att*R with an anticipated size of ∼11.5 Kbp) were identified. Instead, two distinct plasmid populations were recovered, that of pDONR-yUtp2 (Zeo resistant) and p415GPD-3xHA-GW-yUtp2 (amp resistant), despite the fact that the Gateway LR cloning reaction was initially plated on LB/amp (with no Zeo) and that the plasmids were extracted from an overnight LB/amp liquid culture. Thus, these results unexpectedly reveal that the ∼2 Kbp “mystery” band seen during Expression clone *Bsr*GI restriction digestion validation (Fig. 1a) is likely the Donor vector backbone. As well, the Zeocin resistance shown by certain colonies (Fig. 1b) is the result of bacterial plasmid co-transformation of both the p415GPD-3xHA-GW-yUtp2 Expression and pDONR-yUtp2 Entry vectors. According to accepted ideas about plasmid incompatibility and antibiotic selection, the simultaneous transformation and maintenance of these two vectors should be highly unlikely, as both contain the same high copy-number ColE1/pMB1/pBR322/pUC origin of replication (Suppl. Fig. 2) (Velappan et al., 2007). Additionally, there was no selection (Zeo) for the pDONR-yUtp2 Entry vector on the initial transformation plate, as the LR reaction was plated on LB/amp to select for the Expression vector.

To ensure that the pDONR-yUtp2 plasmid solution was not contaminated with Expression vector leading to plasmid co-transformation (despite sequence analysis showing no evidence of this), we performed a transformation of the pDONR-yUtp2 Entry vector (Zeo resistant) into *E. coli* DH5α cells. The transformation was plated on LB/Zeo and LB containing carbenicillin (LB/carb), a more stable analog of ampicillin (in case the co-transformation events were due to breakthrough/satellite colonies from ampicillin) (Sambrook & Russell, 2001). Following overnight incubation, the LB/Zeo plate contained many colonies, while there was no growth on the LB/carb plate, as expected. To check the p415GPD-3xHA-GW-*att*R plasmid solution for contaminating Entry vector, we performed a transformation of the p415GPD-3xHA-GW-*att*R Destination vector into DH5α *E. coli*. The transformation was only plated on LB/Zeo, as any successful transformants (those taking up the Destination vector) would produce the toxic CcdB protein and fail to grow. Therefore, only cells transformed with potential contaminating Entry plasmid would grow. As expected, we saw no growth on the LB/Zeo plate. Additionally, to be certain that carb is equivalent to amp (in that it permits co-transformation to occur), we performed two identical LRs of pDONR-yEmg1 (pDONR-Zeo encoding the *S. cerevisiae* YPH499 NEP1/EMG1 gene) and p415GPD-3xHA-GW-*att*R, plating one on LB/amp and the second on LB/carb. Both plates were then screened for co-transformants by transferring several colonies to LB/amp/Zeo and LB/carb/Zeo plates. Colonies grew on both plates, indicating that co-transformation had taken place, and that co-transformation events are not due to breakthrough growth in the presence of ampicillin.

Based on these unexpected results, we wanted to confirm the presence of two different plasmids in the potentially co-transformed colony using an orthogonal method. For this, we sought to electrophoretically separate the two different plasmid populations by resolving them in the presence and absence of ethidium bromide (EtBr). This dye intercalates DNA, changing the degree of supercoiling of plasmid DNA and thus its migration in a gel (Oppenheim, 1981). This method was historically used to differentiate between linear DNA (where EtBr binding does not change the supercoiling and thus compaction of the molecule due to its free ends) and closed circular DNA (with EtBr binding changing the supercoiling and thus compaction of the plasmid DNA and thus its migration). Here, we used this strategy to confirm that the potentially co-transformed cells carry two different plasmid populations by observing changes in their relative migration when resolved in the presence or absence of EtBr. For this, we resolved the undigested plasmid DNA isolations (the putative Entry/Expression co-transformants, p415GPD-3xHA-GW-yUtp2 and pDONR-yUtp2 (Fig. 1d)) in the presence and absence of EtBr. For the putative Entry/Expression co-transformants resolved in a gel and stained post-run with EtBr (Fig. 1d (i), -EtBr, Mixed Lane), one band at ∼ 7 Kbp is seen co-migrating with p415GPD-3xHA-GW-yUtp2 (Expression Lane) while a second band co-migrates with the pDONR-yUtp2 sample which produced a band at ∼ 4.5 Kbp (Entry Lane).

We repeated the gel in the presence of EtBr, which produced a greater separation of the two plasmid populations within our mixed Entry/Expression sample allowing us to observe two bands at ∼ 10 and ∼ 5.5 Kbp (Fig. 1d (ii), +EtBr, Mixed Lane). Much like the co-migration seen in our initial diagnostic *Bsr*GI restriction digest (Fig. 1a), these bands co-migrated with p415GPD-3xHA-GW-yUtp2 (Expression Lane) and pDONR-yUtp2 (Entry Lane), respectively. As these plasmids have sizes of 9415 and 4509 bp, we found that isolations treated with EtBr prior to and during electrophoresis migrate as slightly larger molecules relative to their expected size. This shift in size of the undigested plasmids between the initial gel (resolved in the absence of EtBr) and this one (resolved in the presence of EtBr) suggests that the bands seen in both gels are in fact plasmid and not contaminating chromosome DNA, as closed plasmid conformations shift in size in the presence of EtBr, but linear DNA does not (Oppenheim, 1981).

In each gel (*Bsr*GI-digested plasmid (Fig. 1a) and undigested plasmid resolved in the presence and absence of EtBr (Fig. 1d)), the presence of bands in the mixed Entry/Expression sample that co-migrate with the Entry and Expression single plasmid bands confirms the identity of the “mystery band” as being (from pDONR) a mixed population of Entry and Expression vectors. This also validates the results of the Oxford Nanopore sequencing by confirming that the mixed population is not an artifact of the sequence assembly and/or contiging process. Additionally, we suspect that the presence of both the Entry and Expression vectors within our mixed Entry/Expression sample is the result of a co-transformation event (*i*.*e*. two plasmids in a single cell) and is not simply a mixed population of cells with different plasmids within the same colony. A study by Tomoiaga et al. (2022) also observed plasmid co-transformation and localized several different plasmids to a single cell through the use of an assortment of fluorescently labeled proteins, demonstrating that a single cell can be simultaneously transformed with two (or more) different plasmids.

### Plasmid co-transformation occurs regardless of Entry and Destination vectors, inserts, antibiotic type, and *E. coli* strain

Having confirmed that the “mystery” band is due to a co-transformation event, we sought to determine whether this occurs with other Gateway vectors. To test this, we performed another LR reaction using a different gene insert (pDONR-yEmg1 rather than pDONR-yUtp2) with p415GPD-3xHA-GW-*att*R, transformed this into *E. coli* DH5α cells, and screened for co-transformants (as described above). Despite using a different gene insert, several co-transformants were observed. Seeing this, we wanted to use both a different Entry and Destination vector with a wider choice of antibiotic resistance to determine if co-transformation would still occur. We performed two LR reactions using the pDONR223-CIRH1A Entry vector (human Cirhin gene) with both pGAD-T7-GW and pGBK-T7-GW Destination yeast two-hybrid plasmids (Lu et al., 2010). The pDONR223 backbone carries a spectinomycin (spec) resistance gene and the pGAD and pGBK backbones contain ampicillin and kanamycin (kan) resistance genes, respectively. Therefore, LB/amp/spec or LB/kan/spec plates exclusively allow the growth of co-transformant colonies. Once each LR had been transformed, plated onto LB/amp or LB/kan, and replica plated onto either LB/amp/spec or LB/kan/spec, we observed several co-transformant colonies on both double antibiotic plates, accounting for ∼9% (pGAD, LB/amp/spec) and ∼29% (pGBK, LB/kan/spec) of transformants. This confirms that co-transformation events are not exclusive to specific Gateway plasmids (and therefore not an artifact of these specific vectors) and can occur while using other plasmids with different combinations of gene inserts and antibiotic resistance genes.

We also wanted to determine if other cloning strains of *E. coli* (beyond DH5α) are amenable to co-transformation. For this, we used the CcdB-resistant DB3.1 *E. coli* strain, as it is used in the Gateway cloning system as a means of propagating and maintaining Donor and Destination vectors. An LR reaction of pDONR-yUtp2 with p415GPD-3xHA-GW-*att*R was performed, transformed into DB3.1 competent cells, and plated on two LB/amp plates before being replica plated onto two LB/Zeo plates. Following transformation, both initial LB/amp plates had hundreds of colonies, with 1255 transformants in total. The LB/Zeo replica plates had 137 transformants in total, equivalent to ∼11% of the colonies on the respective original LB/amp plates. Because these colonies had originally been growing on LB/amp before being replica plated on LB/Zeo, we assume they carry both ampicillin and Zeocin resistance (respectively) and are co-transformants. Therefore, the percentage of co-transformation for these plates at ∼11% is within the range of co-transformation we observe in *E. coli* DH5α (approximately 9-29%). Our results again oppose accepted beliefs about single-plasmid transformation clonality, antibiotic selection and maintenance, and demonstrates that the limitations of transformation are not as rigid as once thought.

### Plasmid co-transformation is not a rare event

Seeing that the phenomenon of co-transformation occurs with different gene inserts and Entry and Expression vectors, with different antibiotics, and in different strains of E. coli, we sought to more precisely determine the frequency of these types of events. During the preliminary screen of eight transformants from the LR reaction of pDONR-yUtp2 and p415GPD-3xHA-GW-*att*R, two of the eight (25%) demonstrated growth on ampicillin and Zeocin (Fig. 1b). This indicated the presence of both the Entry and Expression plasmids and gave an initial estimate of co-transformation frequency.

We expected this number to change when co-transformation frequency was tested using a larger sample size. Therefore, the LR reaction described in the previous section (pDONR-yEmg1 with p415GPD-3xHA-GW-*att*R) was performed with the intention of screening all resulting transformants. Each of the resulting 64 colonies was dashed on LB/amp and LB/amp/Zeo plates. A few colonies (∼12) were too close to one another and could not be cleanly picked. Of the screened colonies, 15 of the 64 transformants (∼23%) showed double antibiotic resistance. To our surprise, this percentage is similar to the 25% rate of co-transformation we saw during the small-scale screening that was initially performed and falls within the 9-29% range of co-transformation described above.

### Unselected plasmids are maintained in co-transformed bacterial colonies

In the course of an LR reaction to create an Expression clone, the recombination reaction is plated on antibiotic media based on the Expression vector’s marker. Despite a lack of antibiotic selection during transformation plating of the pDONR-yUtp2, pDONR-yEmg1, and pDONR223-CIRH1A Entry vectors, all three were co-transformed along with their respective Expression vectors. Based on our understanding of plasmid maintenance and its relationship to antibiotic selection, we had expected these Entry plasmids to be lost after a few passages under non-selective conditions. To investigate how long an Entry plasmid could be maintained in the absence of antibiotic selection, we passaged the 15 co-transformants from the 64 colonies obtained from the LR reaction of pDONR-yEmg1 with p415GPD-3xHA-GW-*att*R described above. Each co-transformant colony was passaged via replica plating or dashing onto new LB/amp plates (for the Expression vector) over the course of seven days in the absence of antibiotic selection for the Entry vector to monitor the maintenance of the unselected Entry vector (Suppl. Fig. 3).

After the first passage, we were surprised to see that all of the co-transformant colonies (screened on LB/amp/Zeo; Suppl. Fig. 3) continued to grow - indicating that no Entry vector plasmids had been lost. Over the seven days that the colonies were passaged (on LB/amp), this trend continued, and none of the initial co-transformants lost their ability to grow on LB/amp/Zeo. After the second and third passages, one of the 15 co-transformed colonies demonstrated poor growth (on LB/amp/Zeo) when compared to the others. However, by the last passage (on LB/amp), this colony was still exhibiting some growth on LB/amp/Zeo. It is unclear if this signalled the beginning of plasmid loss. Interestingly, three other co-transformant colonies exhibited sub-optimal growth (on LB/amp/Zeo) after the fifth and sixth passages (on LB/amp), however, these colonies returned to normal levels of growth on LB/amp/Zeo by the last passage (on LB/amp). These findings are inconsistent with accepted ideas about antibiotics and their requirement in plasmid maintenance, as we expected to see the eventual loss of the unselected pDONR vector. Additionally, the pDONR-yEmg1 and p415GPD-3xHA-GW-yEmg1 vectors share the same type of origin of replication, namely a ColE1/pMB1/pBR322/pUC origin, meaning they are incompatible, and maintenance of the unselected plasmid should be difficult due to the competition for shared replication factors (Sambrook & Russell, 2001; Velappan et al., 2007)

### Plasmid co-transformation is not due to interlocked plasmids

Although sequencing revealed that the LR reaction of pDONR-yUtp2 and p415GDP-3xHA-GW-*att*R did not generate a concatenated plasmid (Fig. 1c (ii)), we wanted to determine whether co-transformation is the result of the creation of interlocking circles during aberrant recombination reactions. For this, we independently generated the p415GPD-3xHA-GW-yUtp2 Expression vector (containing no pDONR-yUtp2 based on the absence of LB/Zeo co-transformants). We then mixed pDONR-yUtp2 and p415GPD-3xHA-GW-yUtp2 based on equal plasmid concentration (∼200 ng/µl) or equal plasmid copy-number. These were transformed into DH5α *E. coli* and plated on LB/amp and LB/amp/Zeo. The LB/amp plates produced a lawn of transformants and the LB/amp/Zeo plates contained thousands of colonies. The growth of colonies on the LB/amp/Zeo plates indicated the presence of both plasmids, suggesting that co-transformation is not a result of the production of interlocking circles during the LR reaction.

### Co-transformation as a consequence of the Gateway cloning system’s experimental set-up

Molecular biology lore, as perpetuated through undergraduate teaching, implies that bacterial plasmid transformation is a single-molecule event, with one plasmid molecule (or population) entering one bacterial cell. The cells then replicate, producing identical copies of the plasmid, and become a clonal colony. Furthermore, it is assumed that one bacterial transformation event (excluding sequential transformations) will very rarely accept more than one plasmid. In rare co-transformation events, the absence of antibiotic selection for one of the plasmids should result in the unselected plasmid being rapidly lost. Plasmid incompatibility suggests that competition for replication machinery and the higher metabolic cost of replication would result in the loss of the unselected plasmid. Our results suggest that these concepts, while still likely true, may not be as absolute as typically presented.

We identified and confirmed through Oxford Nanopore sequencing that the co-transformation of two distinct plasmids, initially p415GPD-3xHA-GW-yUtp2 and pDONR-yUtp2, was occurring during routine Gateway cloning. Further, we found that this phenomenon occurs while using several different plasmid vectors, inserts, antibiotic selection, and strains of *E. coli* and this despite factors meant to prevent co-transformation such as identical origins of replication/plasmid incompatibility and antibiotic selection. We also demonstrated that these co-transformed vectors may be able to stably coexist for longer than once thought in the absence of antibiotic selection. This poses a potential issue for users of the Gateway cloning system, as two (or more) plasmids may be transformed during routine cloning and persist in plasmid DNA isolations etc. where they have the potential to influence experimental findings, such as in sensitive growth and epistatic assays (Carroll and Wong, 2018).

As we have shown, the presence of Entry, Destination, and Expression clones, and of the discarded Gateway vector within a single reaction mixture is characteristic of Gateway cloning and carries with it the risk of co-transformation. We propose that these events are the result of the experimental set-up of a Gateway reaction and not of the recombination itself. In Gateway cloning, equimolar amounts of Donor and Destination vector are mixed. While the unrecombined Destination and the discarded Gateway vectors are eliminated due to the presence of the toxic *ccdB* gene, unrecombined Donor vector remains present at high concentrations in the solution and can co-transform cells, as we have shown. Despite this, Gateway cloning remains a method of choice due to its ease of use, versatility, accuracy, and efficacy, and provides a reliable technique for the movement of DNA inserts between different vectors.

Co-transformation is not seen in conventional cloning. We propose that this is because the plasmid carrying the insert is linearized by restriction digestion (and is thus no longer transformable). The plasmid backbone is often further removed by gel purification of the insert prior to ligation into a new vector (the equivalent of an LR reaction). Thus, any remaining transformable donating plasmid would have resisted restriction enzyme linearization and gel purification and would be of very low concentration.

Based on this, co-transformation is not seen in conventional restriction enzyme and ligation cloning not because it could not occur, but because the plasmid donating its insert is linearized (and thus not transformable) and is purified away. In the case of Gateway cloning, the unrecombined Donor/Entry plasmid remains in a transformable closed-circle configuration and is not purified away. Therefore, the original notion of plasmid co-transformation being a rare event might be the result of the strategy used in conventional restriction enzyme and ligation-based cloning and not a feature of *E. coli* transformation overall.

### Co-transformant screening and validation - experimental recommendations

Here, we propose a simple experimental strategy that we routinely use for screening co-transformant colonies. Following the LR reaction and its subsequent transformation into a cloning *E. coli* strain, 4-10 colonies should be chosen and dashed on LB/amp and LB/amp/Zeo (or the corresponding antibiotic for the desired Expression clone and for the Entry and Expression vector co-transformants) and used to inoculate LB/amp liquid media for plasmid DNA extraction. Any clones growing on LB/amp/Zeo plates (or on media containing the antibiotics for the Entry and Expression vectors) are the result of a co-transformation event and should be discarded. Therefore, successful clones are those growing exclusively on LB/amp (or the Expression vector’s selection antibiotic), making them candidates for plasmid DNA isolation. Diagnostic *Bsr*GI restriction digestion of the plasmid preparation is used to confirm band sizes and that the sample is not a mixed population of Entry and Expression vectors (*i*.*e*., a band corresponding to the size of the Entry vector backbone should not be present).

## Methods

### Antibiotics

Antibiotics were used at the following final concentrations: Ampicillin (amp; 100 µg/ml), carbenicillin (carb; 100 µg/ml), chloramphenicol (chlor; 35 µg/ml), kanamycin (kan; 50 µg/ml), spectinomycin (spec; 100 µg/ml), Zeocin (Zeo; 50 µg/ml).

### Gateway LR cloning

LR reactions were essentially performed as per manufacturer’s instructions using approximately 200 ng/µl of pDONR Entry plasmid (typically 1-3 µl), ∼150 ng/µl of Destination plasmid (1-3 µl), TE to 4 µl, and 1 µl of LR clonase enzyme mix (Invitrogen) for a final reaction volume of 5 µl. The reaction mixture was incubated overnight at room temperature. The next day, 1 µl of Proteinase K was added to the reaction, followed by incubation at 37°C for 10 mins.

### DNA transformations

Plasmid transformations were performed using chemically competent *E. coli* DH5α or DB3.1 CaCl_2_ cells made in-house. To ensure that the competent cells were not already carrying a plasmid, they were plated on 1) Luria broth (LB) and LB supplemented with fresh ampicillin (LB/amp) or Zeocin (LB/Zeo) for DH5α cells, or 2) on LB or LB/amp for DB3.1 cells, then incubated overnight at 37°C. The next day, both the DH5α and DB3.1 cells were growing only on the un-supplemented LB, as expected.

For all transformations, competent cells were thawed on ice prior to the addition of 2 µl of the LR recombination reaction or 1-2 µl prepared plasmid DNA. This was followed by a 20 min incubation on ice and 1.5 min heat shock at 42°C after which the cells were immediately returned to ice. Cells were recovered with the addition of 250 µl LB and incubated with shaking for 1 hour at 37°C prior to plating onto the appropriate antibiotic. Plates were grown overnight at 37°C.

### Initial clone screening

After overnight growth of the transformation plates, several clones were chosen for screening. Each candidate colony was picked using a sterile toothpick and dashed onto new LB/amp and LB/amp/Zeo plates and then used to inoculate 3 ml of LB/amp liquid media. Cultures were grown overnight at 37°C with shaking (180 rpm). The next day, liquid cultures were selected for plasmid extraction based on their ability to grow on LB/amp but not on LB/amp/Zeo plates.

### Plasmid DNA extraction, restriction digestion, and gel electrophoresis

Plasmid DNA extraction was performed using a QIAprep Spin Miniprep Kit (Qiagen) according to the manufacturer’s instructions. For each sample, 2 µl of plasmid DNA was digested overnight at 37°C with the *Bsr*GI restriction endonuclease (*Bsr*GI HF, New England Biolabs) and resolved in a 1.5% agarose/1xTBE gel. The gel was then stained with ethidium bromide (EtBr) and visualized using a Bio-Rad ChemiDoc Imaging System.

### DNA quantification and sequencing

Plasmid DNA was quantified using an Implen NanoPhotometer. For each sample, 600 ng was sequenced by long Oxford Nanopore reads (Plasmidsaurus; Eugene, Oregon, USA).

### Electrophoresis in the presence or absence of ethidium bromide

For each sample, 500 ng of plasmid DNA was loaded per lane in a final volume of 15 µl. Samples without EtBr (-EtBr) were resolved in a low percentage (0.5% agarose/1xTBE) gel. For samples resolved in the presence of EtBr, 0.75 µl (5 ng/ml) of EtBr was added (+EtBr) to the plasmid DNA in a final loaded volume of 15 µl and resolved in a 0.5% agarose/1xTBE gel containing 0.5 µg/ml EtBr. Gels were run overnight for ∼16 hours, were then both stained with EtBr and visualized.

### Replica plating

Replica plating was performed by gently pressing a sterile velvet on a cylindrical block to the plate containing colonies/transformants, then lifting and stamping the velvet onto a new plate.

### Plate photographs

Plate photographs were taken using a Bio-Rad Molecular Imager Gel Doc XR+ on a matte black background.

## Reagents

| Strain | Genotype | Available from |
| --- | --- | --- |
| DB3.1 | F<sup>-</sup> *gyrA462 endA1 Δ(sr1-recA) mcrB mrr hsdS20(r<sub>B</sub><sup>-</sup>, m<sub>B</sub><sup>-</sup>) supE44 ara14 galK2 lacY1 proA2 rpsL20(Sm<sup>R</sup>) xyl5 Δleu mtl1* | Discontinued; equivalent alternative includes the One Shot™ *ccdB* Survival™ 2 T1R Competent Cells from ThermoFisher Scientific |
| DH5α | F<sup>-</sup> *φ80lacZΔM15 Δ(lacZYA-argF)U169 recA1 endA1 hsdR17(r<sub>K</sub><sup>-</sup>, m<sub>K</sub><sup>+</sup>) phoA supE44 λ<sup>-</sup>thi-1 gyrA96 relA1* | Invitrogen/ThermoFisher Scientific |

| Plasmid | Genotype | Description |
| --- | --- | --- |
| p415GPD-3xHA-GW-*attR* | Amp<sup>R</sup>/Cm<sup>R</sup>, LEU2 | Empty yeast overexpression Gateway Destination vector with 3xHA N-terminal fusion. Ref: PMID: 7737504 and 2587792, AddGene: #63781 |
| pDONR/Zeo-yUtp2 | Zeo, yeast UTP2 | Gateway Donor vector with Zeo resistance and a yeast Utp2 insert. Manuscript in preparation. |
| pDONR/Zeo-yEmg1 | Zeo, yeast EMG1 | Gateway Donor vector with Zeo resistance and a yeast Emg1 insert. Manuscript in preparation. |
| pDONR223-CIRH1A | Sm<sup>R</sup>, human CIRH1A | Gateway Donor vector with spec resistance and a human cirhin insert. Available from transOMIC Technologies/Skyang Bio. |
| pGAD-T7-GW-*attR* | Amp<sup>R</sup>/Cm<sup>R</sup>, LEU2 | Empty yeast two-hybrid activation domain/prey Gateway Destination vector with amp and chlor resistances. Ref: PMID: 19843313, AddGene #61702 |
| pGBK-T7-GW-*attR* | Kan<sup>R</sup>/Cm<sup>R</sup>, TRP1 | Empty yeast two-hybrid DNA-binding domain/bait Gateway Destination vector with kan and chlor resistances. Ref: PMID: 19843313, AddGene #61703 |

## Acknowledgements

We thank all members of the Charette Lab for their helpful discussions and critical reading of the manuscript. We thank Maureen Cesmystruk and Sylvia Henry for their administrative support. We thank Haziqa Kassim from the lab of Dr. Meaghan Jones in the Department of Biochemistry and Medical Genetics at the University of Manitoba for their help with Plasmidsaurus sequencing. We also thank Dorothy Zbornak Boudreau-Charette and Eleanor and Theodore Firman-Geer for their editorial assistance.

## Funding

This work was supported by operating grants from the Natural Sciences and Engineering Research Council of Canada (NSERC-DG; RGPIN-2016-06599 and RGPIN-2023-04986), a Dr. James C. Haworth and Dr. Irene Uchida small Genetic/Metabolic grant (SMG2019-03) from the Children’s Hospital Research Institute of Manitoba (CHRIM), a collaborative Canada Foundation for Innovation-John R. Evans Leaders Fund (CFI-JELF) infrastructure grant to J.M.C., and by Brandon University.

## Author Contributions

Courtney L. Geer: conceptualization, investigation, methodology, validation, formal analysis, visualization, writing - original draft. J. Michael Charette: conceptualization, funding acquisition, formal analysis, supervision, writing - review editing.

## Notes

### Competing Interest Statement

The authors have declared no competing interest.

