## Supplemental Figures 1-3. for "An artifact of recombinatorial cloning challenges established beliefs of plasmid co-transformation, selection, and maintenance"

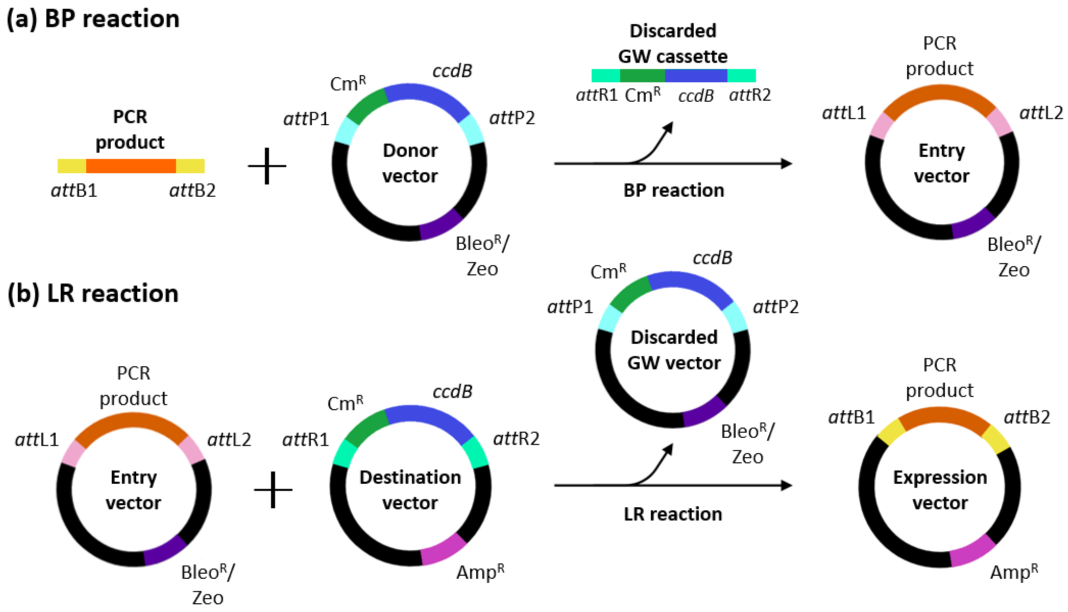


**Supplemental Figure 1. Gateway cloning comprises two recombination reactions.**

**(a)** BP reaction: The *att*B sites (flanking both sides of an amplified PCR product or DNA insert) are recombined with the *att*P sites (flanking both sides of a Gateway cassette within a Donor vector) using Gateway BP clonase enzyme mix to produce *att*L (left) sites. This produces an Entry vector that contains the PCR product (now flanked by *att*L sites) and an excised Gateway cassette which is discarded. Cells transformed with the unrecombined Donor vector are killed by the toxic CcdB protein and thus give no colonies. **(b)** LR reaction: The *att*L sites within the Entry vector are recombined with the *att*R (right) sites of a Destination vector also containing a Gateway cassette to regenerate *att*P and *att*B sites. LR clonase enzyme mix catalyzes this reaction and excises the PCR insert from the Entry vector, then recombines it with the Destination vector. This results in an Expression vector where the PCR insert is flanked on each side by *att*B sites, and a second discarded Gateway vector which includes the toxic *ccdB* cassette. Abbreviations: Amp$^R$ (ampicillin resistance), Bleo$^R$/Zeo (bleomycin/Zeocin resistance, referred to as Zeo; other Donor vectors use kanamycin (Kan$^R$) or spectinomycin (Sm$^R$)), Cm$^R$ (chloramphenicol resistance), and *ccdB* (control of cell death B).

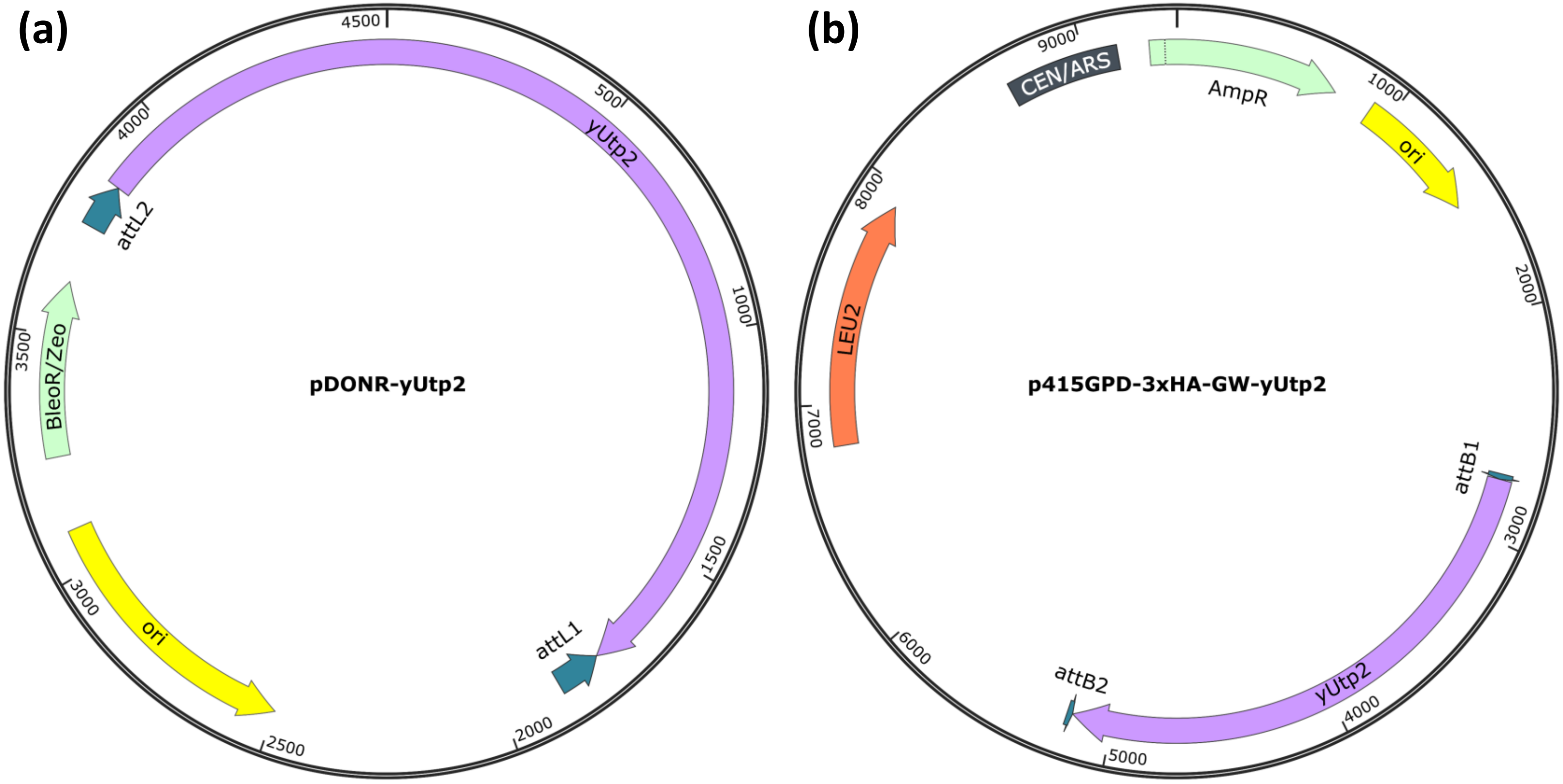


**Supplemental Figure 2. Entry and Expression vectors are sometimes co-transformed during Gateway cloning.**

The two plasmids recovered in the co-transformation events described in Figure 1 include: **(a)** The sequenced 4509 bp pDONR-yUtp2 Entry vector contains a ColE1/pMB1/pBR322/pUC origin of replication (ori), *att*L sequences flanking the yUtp2 gene, and a Zeocin resistance gene, Bleo$^{R}$/Zeo. **(b)** The sequenced 9415 bp p415GPD-3xHA-GW-yUtp2 Expression vector also contains a ColE1/pMB1/pBR322/pUC origin of replication (ori), as well as *att*B sequences flanking the yUtp2 gene insert, an ampicillin resistance gene (Amp$^{R}$), a yeast centromeric origin fused to an autonomously replicating sequence (CEN/ARS), and a LEU2 gene for selection in yeast, along with other features not shown here.

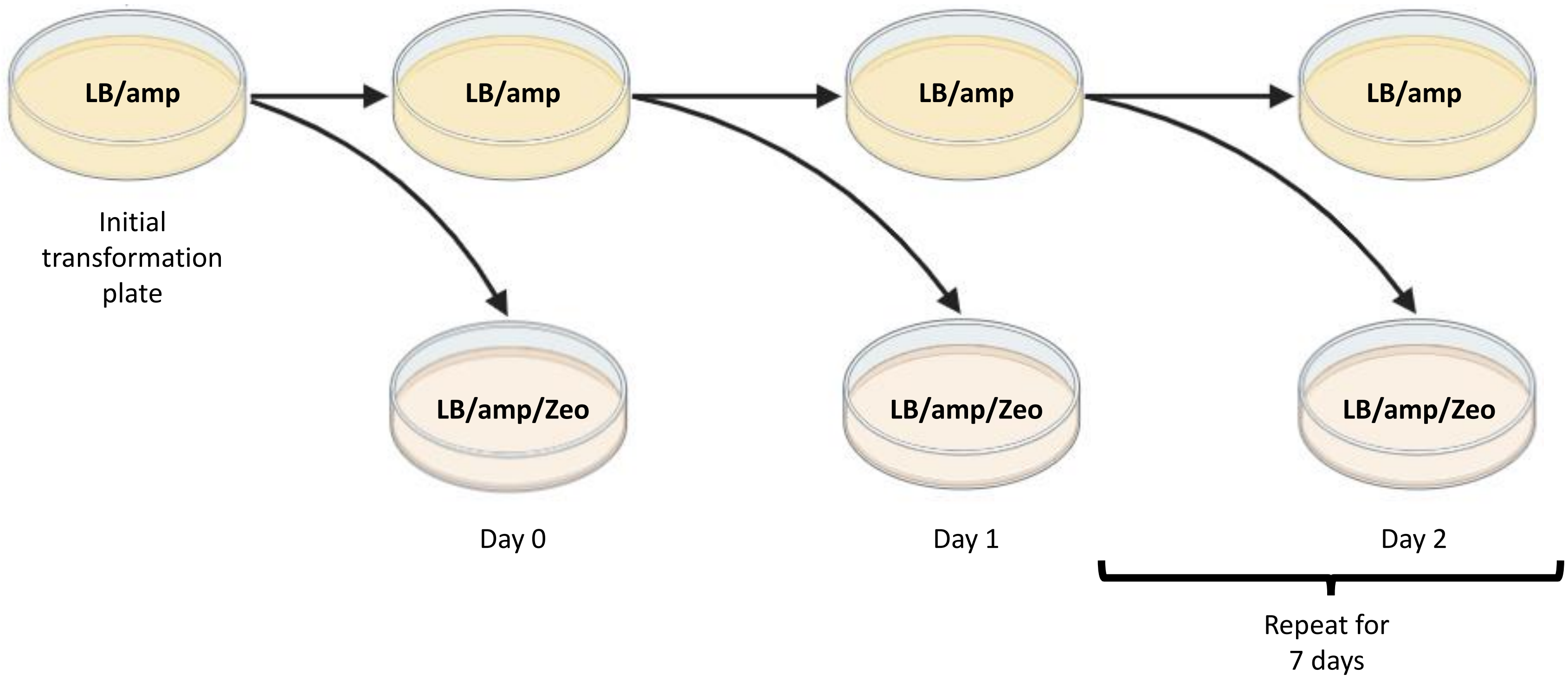


**Supplemental Figure 3. Passaging of transformants to identify and monitor the maintenance of co-transformants.** Each colony on the initial transformation plate was dashed with a toothpick onto an LB/amp plate, then onto an LB/amp/Zeo plate. The plates were incubated at 37°C overnight. For the next seven days, colonies were transferred from the previous day's LB/amp plate (by either replica plating or dashing with a toothpick) to new LB/amp and LB/amp/Zeo plates to monitor plasmid maintenance of pDONR-yUtp2 (Zeo resistant) via the disappearance of colonies over time on LB/amp/Zeo plates.
